# Nuclear morphology predicts cell survival to cisplatin chemotherapy

**DOI:** 10.1101/2022.09.19.508515

**Authors:** Chi-Ju Kim, Anna LK Gonye, Kevin Truskowski, Cheng-Fan Lee, Yoon-Kyoung Cho, Robert H Austin, Kenneth J Pienta, Sarah R Amend

## Abstract

In this study, we characterized nuclear morphology and function as cancer cells underwent recovery following chemotherapeutic treatment to identify the unique characteristics associated with treatment resistance and successful survival. Cells that survived following treatment and resisted therapy-induced cell death were predominantly mononucleated with increased nuclear/cellular size, enabled by continuous endocycling. We found that cells that survive after therapy release likely employ more efficient DNA damage repair and exhibit a distinct nucleolar phenotype - fewer but larger nucleoli – and increased rRNA levels. These data support a paradigm where soon after therapy release, the treated population mostly contains cells with a high level of widespread and catastrophic DNA damage that leads to apoptosis, while the minority of cells that have successful DDR are more likely to access a pro-survival state. These findings suggest that one way cancer cells can survive systemic therapy is to enter the polyaneuploid cancer cell (PACC) state, a recently-described mechanism of therapy resistance. Cancer cells in this state are physically enlarged, undergo whole-genome doubling resulting in polyaneuploid genomes, and are associated with worse prognosis in cancer patients. The PACC state is accessed when a cancer cell experiences external stress, such as genotoxic chemotherapy; after a period of recovery, cells exit the PACC state and resume proliferation to repopulate the tumor cell pool. Our findings demonstrate the fate of cancer cells following chemotherapy treatment and define key characteristics of the resistant PACC state. This work is essential for understanding and, ultimately, targeting, cancer resistance and recurrence.

## Introduction

Once cancer has metastasized, it is incurable because tumors evolve resistance to virtually all systemically administered anticancer therapies. The cells that eventually seed a resistant recurrence must first survive the initial chemotherapeutic insult. While several non-mutually exclusive models exist to explain the predisposition of a cell population to survive therapy (e.g., tumor cell heterogeneity models, cancer stem cell models), less is known about the cell biology characteristics of surviving cells following therapy exposure. Cell morphology is representative of cellular phenotype and reflects the underlying biology of a cell. Abnormal nuclear morphology is often conspicuous in cancer cells [1, 2] and variation in nuclear size and shape is a significant pathologic parameter that is used clinically for the diagnosis of various cancers [1]. Hyper-abnormal nuclear morphologies, such as multilobulated and multinucleated nuclei, are highly associated with metastasis and recurrence [3–5]. Evaluating these cell morphological characteristics is critical to understanding the mechanisms enable cell survival following chemotherapy treatment and the eventual evolution of resistant recurrence.

Chemotherapeutic agents show utility in cancer because of their effects on rapidly proliferating cells, with specific effects on DNA replication machinery or cell division machinery. One underappreciated cellular mechanism to avoid cell death secondary to damage caused by chemotherapy is to skip mitosis and/or cytokinesis, thus not engaging the critical G2/M cell cycle checkpoint. Such mitotic skipping results in increased cellular contents, including increased nuclear size and genomic material, and is hypothesized to aid the cell in adapting to toxic environments such as therapeutic stress [6–9]. The increased genomic content may also help to prevent cells from undergoing cell cycle checkpoint-activated apoptosis due to DNA damage [4, 6–15]. This amitotic cell cycle that leads to genomic duplication is accompanied by abnormal nuclear morphologies including large mononucleated, mononucleated and simultaneously lobulated, or multinucleated phenotypes [6–9, 12–14].

DNA damage response (DDR) is a pathway to overcome the effect of chemotherapeutic agents such as cisplatin, a DNA poison. If cells are exposed to genotoxic stress, DDR machinery is activated to repair damaged DNA. In case of double-stranded DNA breaks, there are two DDR pathways including non-homologous end-joining (NHEJ) and homologous recombination (HR). DDR procedures commonly consist of two crucial steps: sensing damaged DNA lesions and recruiting DNA repair molecules. Gamma H2A.X (gH2AX) acts as an DNA damage sensing marker, and 53BP1 in NHEJ or BRCA1 in HR are active DNA repair molecules. These DDR machineries thus initially allow the cells to survive after chemotherapy treatment. Furthermore, successfully completed DDR may also prevent cell death at cell cycle checkpoint that enables reactivate cell cycles to prolong their cellular life.

To define the life history of cells following chemotherapy treatment that continue to survive versus those that die after chemotherapy, we characterized nuclear morphology and function over time following release from treatment with cisplatin. We demonstrate that mononucleated, polyploid cells dominate the population of cells that survive following therapy and thus likely represent the cells that will eventually seed a therapeutically resistant population. These data indicate that cells that survive cisplatin treatment access a mononucleated state, continue to replicate DNA via endocycling, and retain high levels of DNA damage repair, while multinucleated cells eventually undergo cell death.

## Materials and methods

### Cell culture

PC3 and MDA-MB-231 cell lines were originally purchased from ATCC (United States). Cells were maintained in RPMI-1640 with L-Glutamine (PC3; Gibco, #11875-093) or DMEM (MDA-MB-231 cells; Gibco, #11995-065), with 10% FBS (Avantar, #97068-085) and 1% penicillin streptomycin (Gibco, #15140-122).

### Treatment

Sub-confluent cultures of PC3 cells were treated cisplatin (6 μM; Millipore Sigma, #232120-50MG), docetaxel (5 nM; Cell Signaling Technology, #9886), or etoposide (25 μM; Cell Signaling Technology, #2200S) and cultured for 72 hours under standard conditions. MDA-MB-231 cells were treated with cisplatin at 12 μM. Large cells were then sterilely isolated via size-restricted filtration with a pluriStrainer^®^15 μm cell strainer (pluriSelect, #45-50015-03) and replated for culturing.

### Nuclear morphological analysis

Large cells were generated as described, isolated, and then cultured for 1, 5, 10, and 15 days following 72 hours of drug treatment. 24 hours prior to each timepoint, the cells were seeded into 35 mm IbiTreat *μ*-Dishes (Ibidi USA Inc., #81156) and incubated overnight. Samples were fixed in 10% formalin for 20 minutes, and permeabilized for 10 minutes in 0.2% Triton-X-100 in PBS. Permeabilized samples were blocked with 10% goat serum in PBS at room temperature for 30 minutes. Samples were then incubated overnight at 4°C with a lamin A/C primary antibody and secondary antibody (Table S1) for 1 hour at room temperature. Samples were mounted with ProLong™ Diamond Antifade Mountant with DAPI (Invitrogen, #P36971) and images were acquired using a Zeiss Observer Z1 microscope/ZEN pro 2.0 software (Carl Zeiss Microscopy).

Nuclear morphology (mono-nucleated vs. multi-nucleated) was manually discriminated and counted using Fiji [38].

### Nucleolar morphological analysis

Cells were induced, seeded, fixed, permeabilized, and blocked as above. The samples were incubated with nucleolin and lamin A/C primary antibodies overnight at 4°C and a secondary antibody (Table S1) for 1 hour at room temperature, protected from light. Samples were mounted and images were acquired as described above. Images were analyzed for nucleolar number and area via a custom CellProfiler 4.0 [39] pipeline.

### DNA damage response marker foci analysis

Treated cells were induced, seeded, fixed, permeabilized, and blocked as above. Samples were incubated overnight at 4°C with gamma H2A.X, 53BP1, and lamin A/C primary antibodies (Table S1), followed by washes with 0.1% PBS-T. Appropriate secondary antibodies (Table S1) were incubated for 1 hour at room temperature, followed by five washes with 0.1% PBS-T. Antibodies were diluted with 1% (v/v) bovine serum albumin (BSA, Millipore Sigma, A9418) in PBS. Samples were mounted using ProLong™ Diamond Antifade Mountant with DAPI. Image acquisition was performed using a Zeiss Observer Z1 microscope/ZEN pro 2.0 software (Carl Zeiss Microscopy). Images were analyzed via a custom CellProfiler 4.0 pipeline.

### 3D, confocal, live-cell imaging

Live cells were incubated for 45 minutes at standard conditions with 1 μg/mL of CellTracker™ Orange CMTMR (Invitrogen, #C2927) and 10 μM Vybrant™ DyeCycle™ Green Stain (Molecular Probes, # V35004) to label the cytoplasm and nucleus, respectively. Labeled cells were lifted and resuspended in cold Matrigel Basement Matrix (Corning, #356232), plated onto 35 mm Ibidi *μ*-Dishes (glass bottom, Ibidi USA Inc., #81158), and immediately coverslipped. Plated samples were cultured for 30 minutes at standard conditions to allow for solidification of the Matrigel, and then covered with phenol-free RPMI-1640 with L-Glutamine (Gibco, #1835030) with 10% FBS and 1% penicillin streptomycin. Image acquisition was performed using a Nikon Eclipse Ti2 microscope with a Nikon C2 camera and NIS Elements version 5.11. (Nikon Inc.) with a live-cell imaging ThermoBox incubation system (Tokai Hit Co.). 3D image processing and analysis was performed with Imaris 9.8 software (Oxford Instruments).

### Time-lapse imaging of PC3-FUCCI

PC3-FUCCI cells were plated and cultured overnight under standard conditions (see above) and then treated with 6 μM cisplatin for 72 hours. After treatment, media was changed and live-cell timelapse microscopy of treated cells was performed in an Incucyte SX5 (Sartorius). Images were taken every 30 minutes in the phase, green, and orange channels.

### Flow cytometry analysis

Live cells were incubated with a FITC-Annexin V (BioLegend, #640906) according to the manufacturer’s recommendations. Cells were analyzed on an Attune NxT Flow Cytometer (Thermo Fisher Scientific). At least 40,000 events were recorded per sample and acquired data was analyzed using FlowJo software version 10.8 (BD Biosciences).

### Immunofluorescent imaging of autophagosomes

Cells were seeded into 35 mm IbiTreat *μ*-Dishes and cultured overnight. The cells were fixed and permeabilized as above. Following permeabilization, samples were blocked with 10% BSA for 30 minutes. The samples were then incubated overnight at 4°C with LC3 A/B, lamin A/C, gamma H2A.X primary antibodies (Table S1) and subsequently washed with 0.1% (v/v) PBS-T. Samples were incubated for 1 hour at room temperature with appropriate secondary antibodies (Table S1), followed by washes with PBS-T. Samples were mounted with ProLongTM Diamond Antifade Mountant with DAPI (see above). Images were acquired using a Zeiss Observer Z1 microscope and ZEN pro 2.0 software (Carl Zeiss Microscopy).

### Autophagosome detection by 3D live-cell imaging

Live cells were labeled with 2 μM Calcein AM (Invitrogen, #C3100MP) for cytoplasm and Autophagy Assay Kit (Red) (Abcam, #ab270790) for 30 minutes under standard culture conditions as recommended by the manufacturer. Labeled cells were washed with HBSS, and phenol red-free media was added to culture dishes prior to imaging. Image acquisition was performed using a Nikon Eclipse Ti2 microscope with a Nikon C2 camera and NIS Elements version 5.11. (Nikon Inc.) with a live-cell imaging ThermoBox incubation system (Tokai Hit Co.). Image processing and analysis was performed with Imaris 9.8 software (Oxford Instruments).

### Statistical analysis

All statistical analyses were performed in GraphPad Prism version 9.1 (GraphPad Software, LLC). An α = 0.05 (confidence level 95%) was the criterion considered to determine statistical significance for all tests. No significance (ns), *, **, ***, and **** represent p values of ≥0.05, <0.05, <0.01, <0.001, and <0.0001, respectively. For violin plots, red lines indicate median and blue lines indicate lower and upper quartiles.

## Results

### Cell death and survival after cisplatin treatment

Prostate cancer cell line PC3 was treated with 6 μM cisplatin for 72 hours. Treatment effect, e.g., a lethal dose (LD) 50, is typically evaluated immediately following therapy exposure, and we observed the expected cell die-off at 72 hours. To characterize the cells that survive cisplatin treatment and then have the potential to go on to seed a resistant recurrence, we removed the chemotherapy and permitted the cultures to recover in complete media. Following release of cisplatin treatment, cells continued to die, though this cell death gradually decreased with time, and eventually plateaued (**Fig. 1A, B**). The number of surviving cells continues to decrease over time, with the highest level of cell death observed in the first three days post-cisplatin release (**Figure 1A, B**). We evaluated apoptosis by flow cytometry for Annexin V stain. We found that the level of apoptosis was higher in the treated cultures compared to untreated control, but this difference decreased with increased recovery time out to three days post-treatment (**Fig. 1C**). To understand the cell biological phenotypes of cells that survive cisplatin treatment, we evaluated key morphological and functional features of surviving cells at various timepoints post-cisplatin treatment.

**Figure 1.**
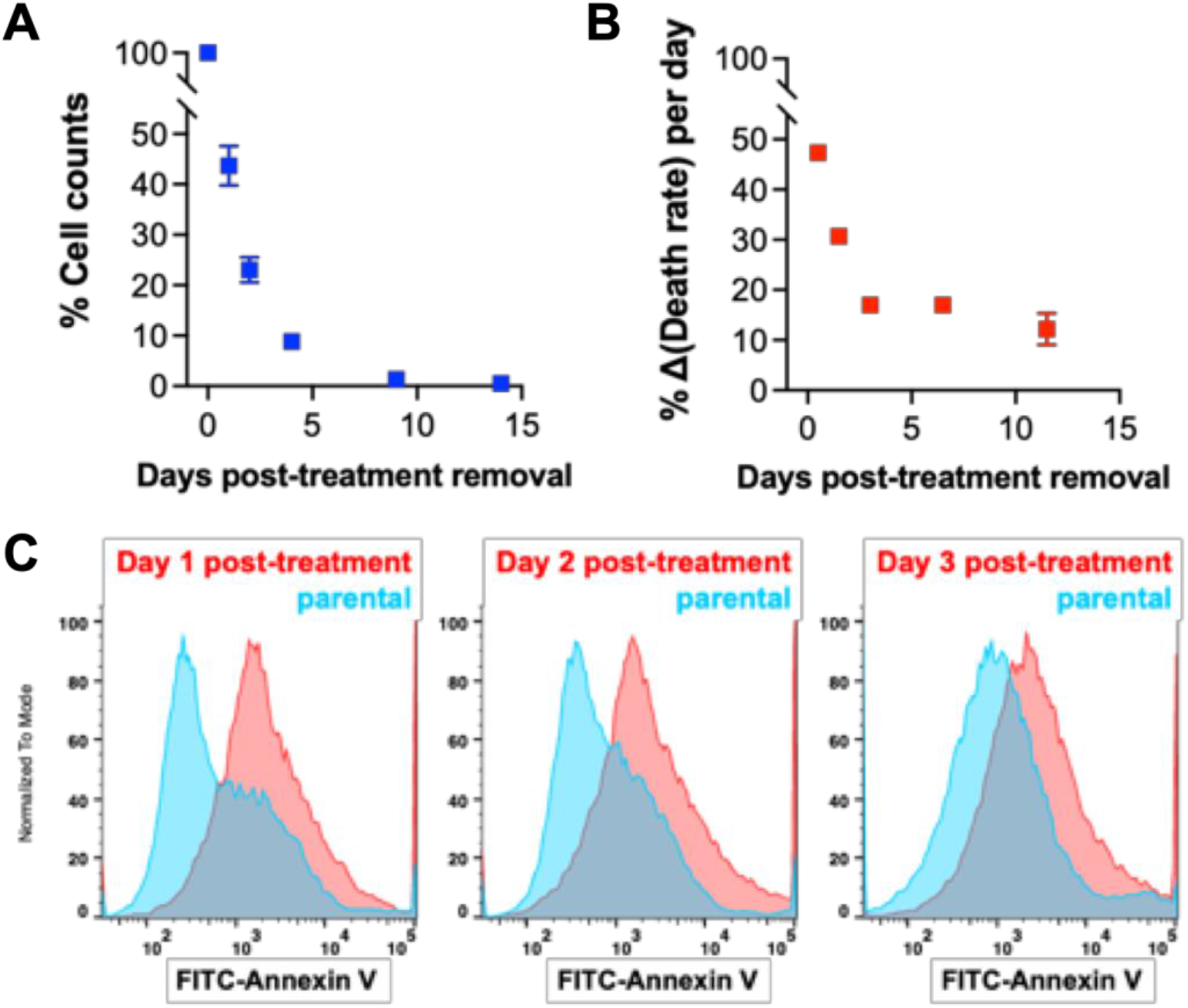
Cell death and survival after cisplatin treatment. Relative cell number (percent normalized to population count at 0 days post-treatment removal) as a function of recovery time **(A)**was determined for treated PC3 cells generated with 72h 6 μM [LD50] cisplatin treatment. From this data, we also determined **(B)**the percent change in the death rate for each of the recovery days we investigated. **(C)**Flow cytometry was used to measure surface expression of annexin V as a marker of apoptosis in treated PC3s generated with 72h [LD50] cisplatin treatment at days 1, 2, and 3 post-treatment removal compared to parental PC3 prostate cancer cells.

### Increasing cellular and nuclear volume and repeated whole genome doubling in surviving cells over time

We assessed cell volume and nuclear volume of treated cells post treatment-release utilizing 3D confocal microscopy. The total cell volume of surviving cells increased over recovery time (**Fig. 2A**). Surviving cells also increased in nuclear volume over time (**Fig. 2B**). The nucleus-to-cytoplasm ratio gradually decreased as a function of recovery time (**Fig. 2C**). This implies that the rate of synthesis of non-nuclear biomolecules is relatively faster than genomic replication compared to treatment-naïve cells. To determine the extent to which the nuclear size was influenced by cytoskeletal/cytoplasmic components, we measured the area of isolated nuclei stained for lamin A/C (**Fig. 2D, E**). Consistent with the whole-cell data, the size of isolated nuclei increased with time from treatment (**Fig. 2E**).

**Figure 2.**
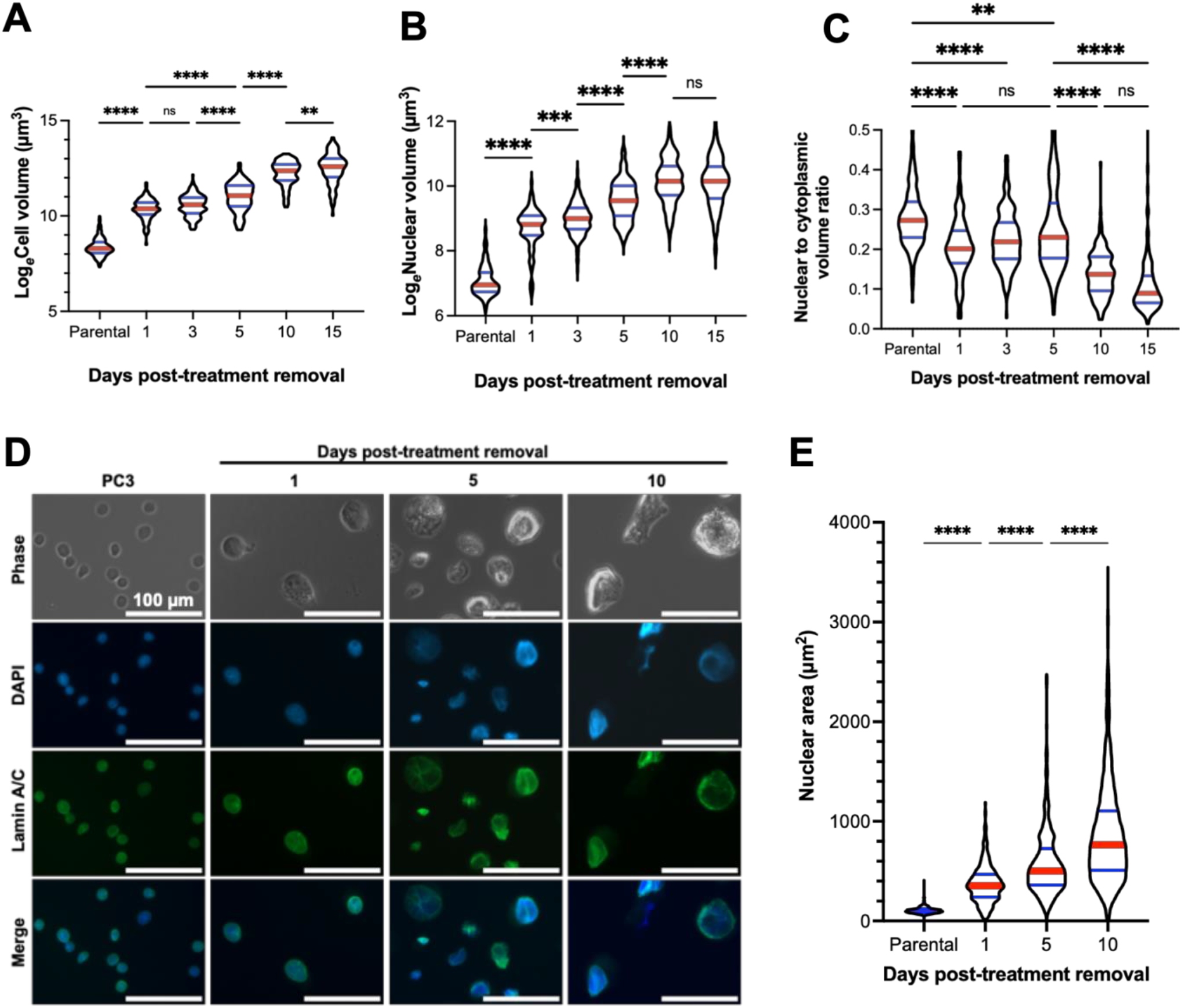
Increasing cellular and nuclear volume in surviving cells over time. Natural log-transformed **(A)**nuclear volume and **(B)**cellular volume of live treated PC3 cells, generated with 72h 6 μM cisplatin treatment, at various timepoints in recovery was determined by suspending cells in Matrigel and staining DNA with Vybrant™ DyeCycle™ Green Stain (for DNA) and CellTracker™ Orange (for cytoplasm). 3D images were obtained via confocal microscopy and images were rendered for volumetric determination using Imaris 9.8 software. Data was analyzed for differences between groups using a one-way ANOVA, a post-hoc Tukey’s multiple comparisons tests was used to detect between-group differences. **(C)**Nuclear and cytoplasmic volumes were used to calculate a nuclear to cytoplasmic volume ratio of treated PC3s at various timepoints past treatment removal. A Kruskal-Wallis test was used to analyze if there were any significant differences among the samples, Dunn’s multiple comparison’s test used to test for comparisons between groups. **(D)**Nuclei were isolated from parental and treated PC3 cells before mounting on PLL-coated dishes. Nuclei were stained for lamin A/C and covered with ProLong™ Diamond Antifade Mountant with DAPI before imaging on a Zeiss Observer Z1 microscope. The size of isolated nuclei was quantitated **(E)**via a custom CellProfiler 4.0 pipeline. A Kruskal-Wallis test was used to analyze if there were any significant differences among the samples, Dunn’s multiple comparison’s test used to test for comparisons between groups. For all tests: * = p<0.05, ** = p<0.01, *** = p<0.001, **** = p<0.0001.

The increasing nuclear size of the surviving cells suggested that mononucleated cells undergo repeated DNA duplication during the recovery period. We utilized a fluorescence ubiquitination cell cycle indicator (FUCCI) cell cycle sensor to track progression through cell cycle (**Fig. 3A**) [16]. The nuclei of cells transduced with the FUCCI construct fluoresce red in G1, yellow (double positive for red and green) in very early S, green in mid-late S, G2, and up to anaphase in M, and are colorless in late-M and G0 (**Fig. 3B**). Using live cell imaging, we observed large cisplatin-treated cells progressing through these different stages without undergoing cell division, a process termed endocycling (**Fig. 2C**). To specifically test for DNA replication, cultures were pulsed with thymidine analogue EdU. Positive EdU staining confirmed that mononucleated cells actively engaged in *de novo* DNA synthesis at all timepoints following removal of cisplatin (**Fig. 3D**). We additionally confirmed that the DNA content in the nuclei of the cells increased during the recovery period by measuring an integrated intensity of DAPI, which also increased over the recovery period (**Fig. 3E**). These findings indicate that surviving cells, while not proliferative, are functionally active in genome duplication.

**Figure 3.**
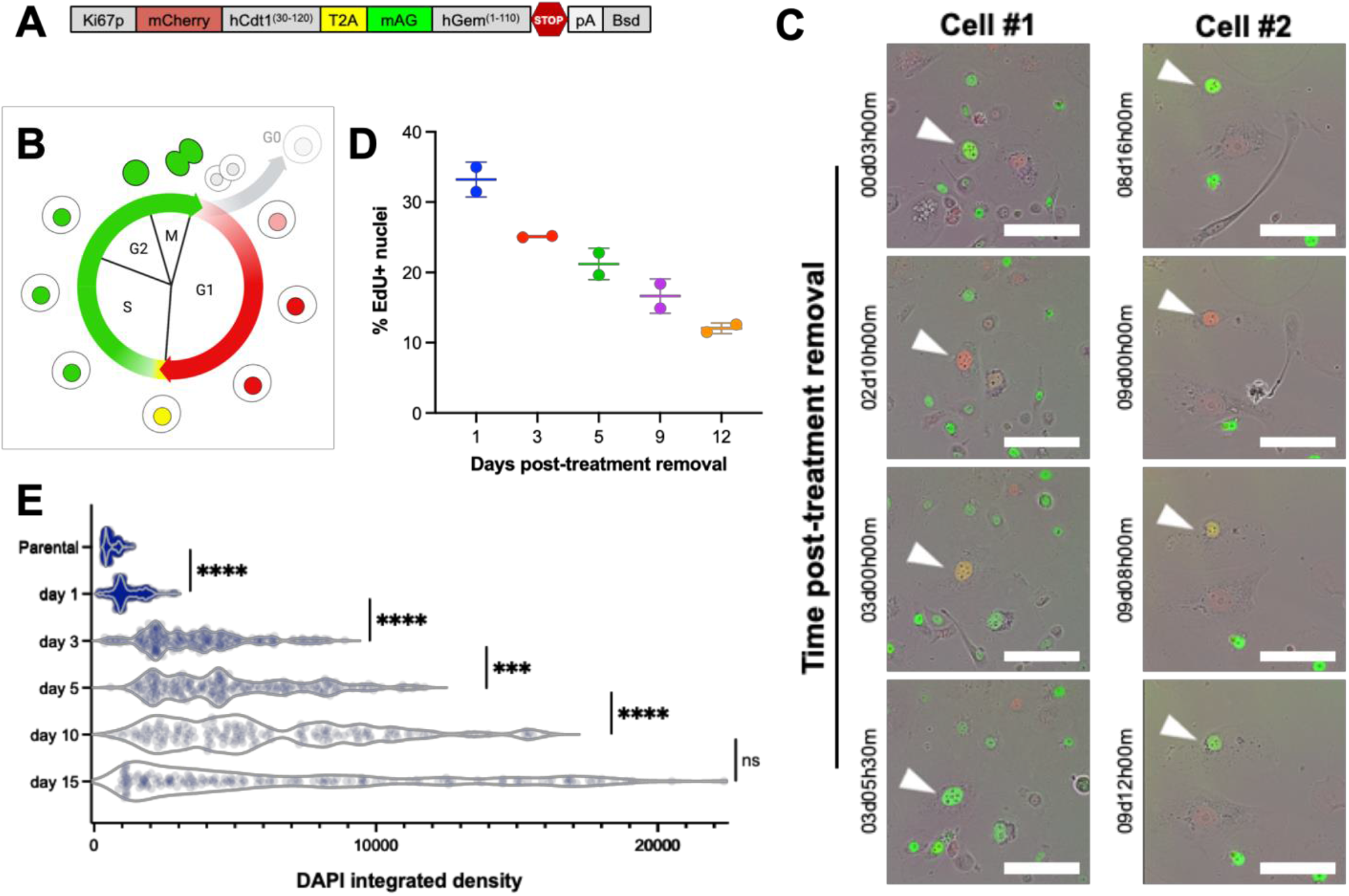
Repeated whole genome doubling in the absence of cell division. PC3 cells were transduced with a **(A)**Ki67p-T2A-FUCCI construct, gifted from Alexander Zambon. **(B, C)**Treated PC3s were generated from this reporter cell line with 72h 6 μM cisplatin treatment, and timelapse microscopy was performed with an Incucyte SX5 with images taken every 30min after release from treatment, scale bar = 200 μm. Two representative cells are shown. **(D)**Cisplatin-treated PC3s were tested for *de novo* DNA synthesis via a Click-It Plus EdU assay according to the manufacturer’s protocol. Images were acquired using the 10x objective on a Zeiss Observer Z1 microscope. A custom CellProfiler pipeline was developed and used to quantify DAPI+Edu+ nuclei. **(E)**PC3 cells were treated with 6 μM cisplatin for 72h, and immunofluorescence imaging was used to visualize genomic contents as a measure of integrated DAPI intensity. This was analyzed for differences between groups using a one-way ANOVA, a post-hoc Games-Howell’s multiple comparisons tests was used to detect between-group differences. For these tests: *** = p<0.001, **** = p<0.0001.

### Nuclear morphology of cells that survive cisplatin treatment

We classified the morphology of treated cancer cell nuclei as mononucleated or multinucleated. Conventional nuclear visualization by DNA staining with DAPI or Hoechst alone is not always suitable for distinguishing nucleus morphology (**Fig. S1**). The nuclear envelope of cells was visualized by performing immunofluorescent labeling of lamin A/C to enhance visualization (**Fig. 4A**). To overcome limitations in 2D methodology due to a spatially hindering effect of the nuclear envelope signal (**Fig. S2**), we also utilized 3D confocal imaging (**Fig. 4B**). Immediately after removal from treatment, treated PC3 cells had a uniformly distributed nuclear morphology, with approximately equal proportion of mono- and multinucleates (**Fig. 4C**). As cells recovered, mononucleates became predominant over time by 2D imaging (**Fig. 4C**) and confirmed by 3D confocal microscopy imaging (**Fig. 4D**). Morphologically, the sphericity of nuclei gradually decreased over recovery time (**Fig. 4G, H**), suggesting that nuclei at later time points may exhibit high deformability, perhaps due to a lower supporting force derived from their cytoskeleton [5, 17–19].

**Figure 4.**
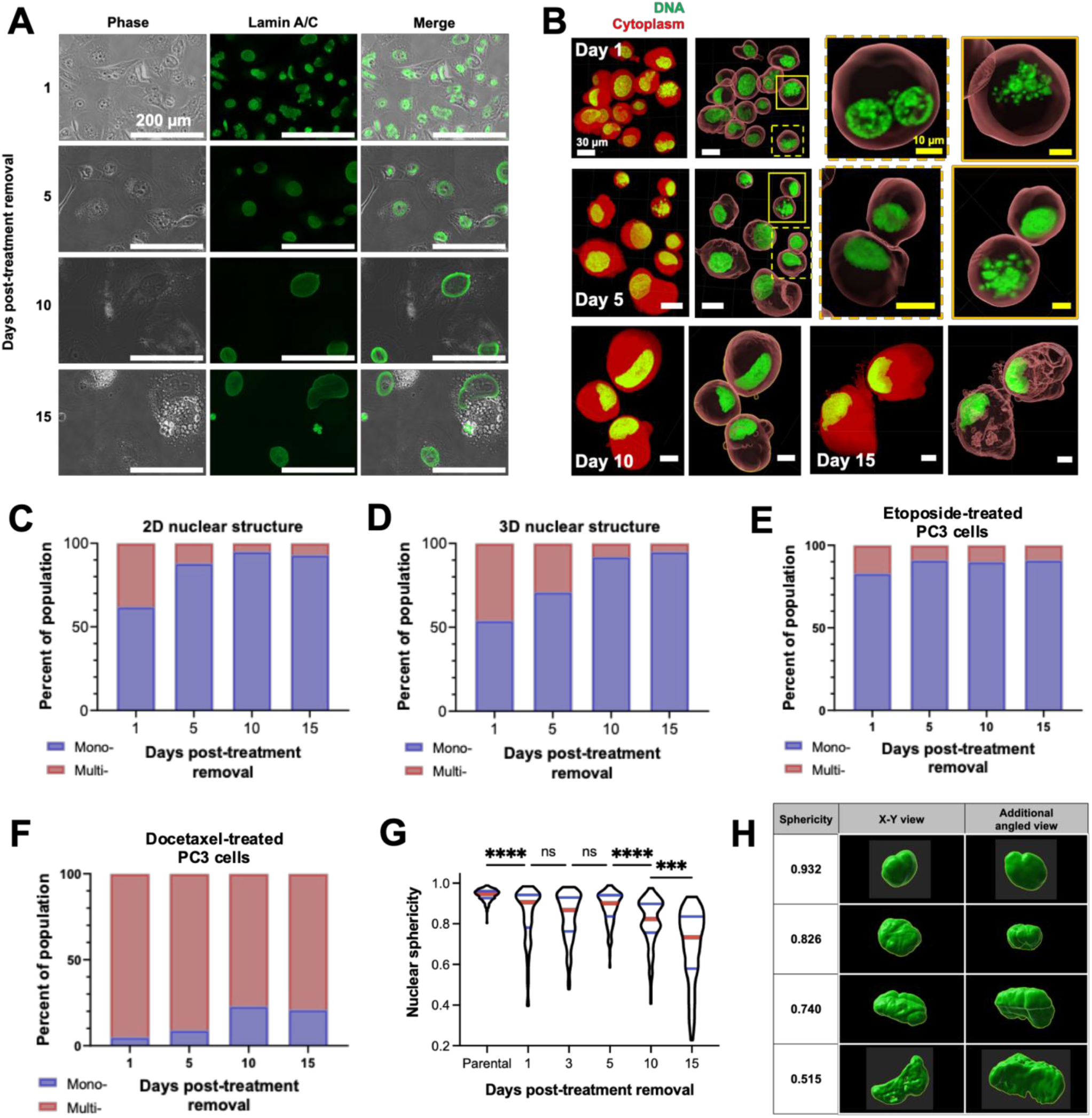
Characterization of the nuclear morphology of cisplatin-treated prostate cancer cells. Representative immunofluorescence images **(A)** to demonstrate the nuclear morphology of treated PC3 cells generated with 72h 6 μM cisplatin treatment following removal from chemotherapy. **(B)** Live treated PC3 cells at various timepoints in recovery were suspended in Matrigel and staining DNA with Vybrant™ DyeCycle™ Green Stain (for DNA) and CellTracker™ Orange (for cytoplasm). 3D images were obtained via confocal microscopy and images were rendered and volumes determined using Imaris 9.8 software. **(C)** Population distribution of mononucleated and multinucleated PC3s, derived from images described in (A). Nuclear morphology (mononucleated vs. multinucleated) was manually discriminated and counted using Fiji. At least 250 cells were analyzed per condition. **(D)** Nuclear morphology was also determined using the 3D images described in (B). The population distribution of mononucleated and multinucleated treated PC3 cells generated with 72h **(E)** 25 μM etoposide treatment or **(F)** 5 nM docetaxel treatment was also determined, derived from immunofluorescent images of cells stained for lamin A/C and mounted with DAPI. Nuclear morphology (mononucleated vs. multinucleated) was discriminated and counted in Fiji software.

To evaluate the drug-specific phenotypic responses to treatment and recovery we evaluated surviving cells following treatment with the common chemotherapeutic drugs etoposide (topoisomerase II inhibitor) and docetaxel (microtubule stabilizer) for 72 hours (**Fig. 4E, F**). The proportion of multinucleates was significantly different between the two drugs and compared to cisplatin. Increased proportion of multinucleated cells in the docetaxel treatment group are likely due to the mechanism of this therapy, which stabilizes microtubules, interrupting mitosis and inhibiting cell proliferation. Regardless of treatment type, all groups showed an increased proportion of mononucleated cells with recovery time. These results suggest that mononucleated PC3 cells will eventually become a dominant morphological phenotype in the longer recovery period.

### DNA damage repair is retained in surviving cells

The DDR plays a crucial role in avoiding programmed cell death and prolonging cellular life [20]. gH2AX foci mark sites of DNA damage, and 53BP1 molecules form distinct foci when recruited to sites of DNA damage to initiate DNA repair [20]. To evaluate the extent of DNA damage and initiation of DDR, expression and colocalization of gH2AX and 53BP1 were assessed for each recovery time point (**Fig. 5A, D**).

**Figure 5.**
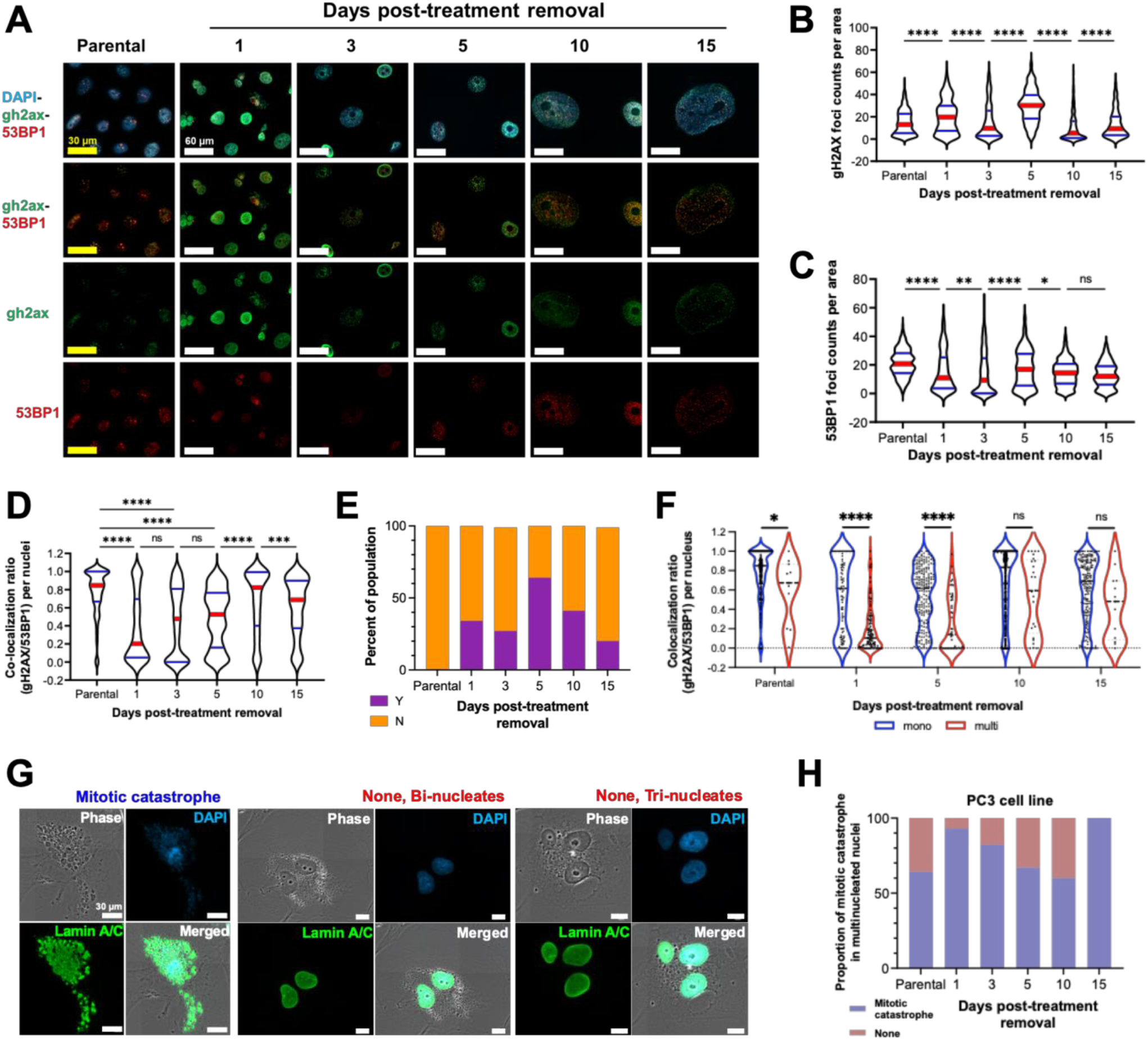
The DNA damage response in cisplatin-treated prostate cancer cells. PC3 cells were treated with 6 μM cisplatin for 72h. **(A)**Immunofluorescence staining and imaging was used to visualize genomic contents (DAPI), sites of DNA damage (gH2AX), and DDR machinery recruitment (53BP1). From these images, a custom CellProfiler 4.0 pipeline was used to determine raw count per unit cellular area of the number of **(B)**gH2AX foci and **(C)**53BP1 foci, as well as **(D)**the colocalization ratio of gH2AX with 53BP1. **(E)**The relative proportion of cells with (Y) and without (N) gH2AX pan-stained nuclei from was determined. **(F)** For each day post-treatment recovery, cells were classified as mononucleates and multinucleates, and the colocalization ratio of gH2AX with 53BP1 was determined for each cell class. Mononucleated, multinucleated, and mitotically catastrophic cells were classified via a custom CellProfiler 4.0 pipeline, and **(G-H)**the proportion of multinucleated cells to those experiencing mitotic catastrophe were plotted across the different days post-treatment removal. Mann-Whitney tests were used to detect differences between groups in (B), (C), (D), and (F). For all tests: * = p<0.05, ** = p<0.01, *** = p<0.001, **** = p<0.0001.

Excessive DNA damage that induces acute apoptosis is distinguished by a pan-staining pattern of gH2AX [21, 22]. The proportion of gH2AX pan-stained nuclei in treated cells was approximately 50% of the total population on day 1 and 30% on day 3 after cells were removed from treatment, gradually decreasing during recovery (**Fig. 5E**); this was consistent with our observations of overall cell death (**Fig. 1A, B**). We also evaluated gH2AX and 53BP1 foci per unit nuclei area in cells whose nuclei did not exhibit pan-stained gH2AX (**Fig. 5B, C**). In these experiments, we observed that the total number of gH2AX foci generally increased early in the period of recovery from treatment, and later decreased to parental levels at days 10 and 15 post-treatment removal. The number of 53BP1 foci decreased until day 3 and then returned to pre-treatment levels by the later recovery timepoints (**Fig. 5C**). Colocalization of gH2AX and 53BP1 foci are indicative of intact DNA damage repair. Early in recovery (days 1 and 3), treated cells had a low level of colocalization, but the colocalized gH2AX/53BP1 foci returned to the level of the untreated cancer cell control at day 10 post-treatment (**Fig. 5D**). Over time, the size of gH2AX foci decreased, whereas the size of 53BP1 foci increased following therapy from day 1 to day 10 (**Fig. S3**). Together, these findings suggest that surviving cells activate the DDR pathway to avoid cell death [20].

We found that mononucleate cells became the dominant population over the duration of treated cell recovery (**Fig. 4C-D**), suggesting that multinucleated cancer cells died. To investigate the activation of the DDR in these distinct nuclear morphologies, the level of colocalization between gH2AX and 53BP1 foci in each type was compared (**Fig. 5F, Fig. S4**). Mononucleate cells contain more colocalized DDR foci than multinucleates, suggesting that mononucleated cells retained intact DNA damage of repair that was not observed in multinucleated cells.

A primary mechanism of multinucleate formation is a mitotic catastrophe [6, 9, 13]. The proportion of nuclei that underwent a mitotic catastrophe in cisplatin treated PC3 was determined by examining immunofluorescent-labeled images of the nuclear envelope (**Fig. 5G, H**). The results indicate that proportion of mitotic catastrophe in treated PC3 cells decreased in the recovery period. These findings suggest that an active DDR suppresses multinucleation due to mitotic catastrophe, preventing cell death.

### Nucleolar size increases and number decreases following cisplatin treatment release

The primary substructure of the nucleus is the nucleolus, an rRNA production hub that plays a significant role in nuclear homeostasis [23, 24]. We found that the number of nucleoli within the nuclei of treated PC3 cells decreased (**Fig. 6A, B**), while the average size of each nucleolus increased (**Fig. 6A, C**) over recovery time. The nucleolus to the nucleus area ratio in treated cells initially decreased at early time points (until day 3) and rebounded during recovery (**Fig. 6D**). These results indicate that the synthesis of nucleolar molecules was initially slower than nuclear molecules in treated cells; however, this area ratio returns to relative levels similar to untreated controls, suggesting the potential of nucleoli to actively respond to genomic stress during treatment recovery to maintain nuclear homeostasis [24, 25].

**Figure 6.**
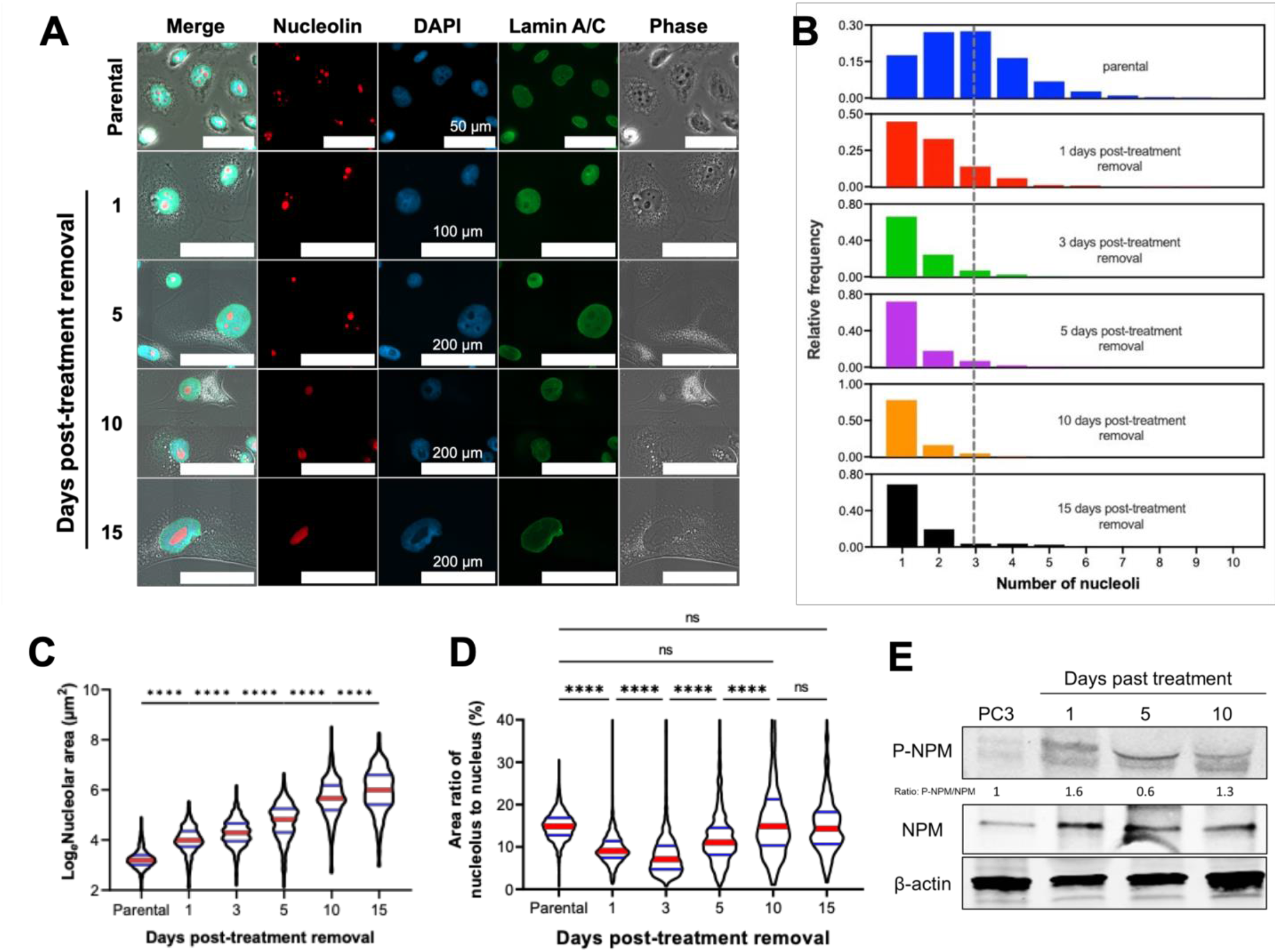
Nucleolar counts, size, and molecular signatures. Treated PC3 cancer cells generated with 72h 6 μM cisplatin treatment were stained for lamin A/C and nucleolin, and were mounted with DAPI, **(A)**representative images of nucleoli are shown. A custom CellProfiler 4.0 pipeline was used to then determine the **(B)**number and **(C)**area of the nucleoli of the treated PC3s throughout the recovery period to become PACCs. Nucleolar area was analyzed for differences between groups using a one-way ANOVA, and a post-hoc Tukey’s multiple comparisons test was used to detect between-group differences. The lamin A/C stain was used to calculate nuclear area, and thus **(D)**the nucleolar to nuclear area ratio at each timepoint posttreatment removal could be calculated; a Kruskal-Wallis test was used to analyze if there were any significant differences among the samples, with a Dunn’s multiple comparison’s post-hoc test. For all statistical tests: * = p<0.05, ** = p<0.01, *** = p<0.001, **** = p<0.0001.

In its function as a ribosomal RNA production hub, a nucleolus contains genes for rRNAs [23, 24]. We found that the level of 28S rRNA in treated PC3 cells was higher than that in non-treated cells and that it increased with recovery time. 18S rRNA expression was lower in treated cells overall, while the level of 5.8S rRNA remained similar in all conditions (**Fig. S5**). The altered rRNA composition of cells recovering from cisplatin treatment suggests that the remodeling of rRNA configuration in the nucleolus may be a response to genomic stress.

## Discussion and conclusions

The majority of studies that describe lethal or inhibitory concentrations of therapies look at a single time point, e.g., how many cells dies at 72 hours. Furthermore, most of the assays used to assess therapeutic effect do not look at the morphology of cells at the time of assay. In this study, we explored the life history of cells in the days and weeks after cisplatin treatment to gain insights into the adaptive characteristics of surviving cells that are likely to seed eventual recurrence. We found that cells that resist therapy-induced cell death are predominantly mononucleated and undergo multiple rounds of genome duplication without cell division in a process of endocycling, resulting in increased nuclear size and increased cell size. We further find that cells that survive after therapy release retain continued DDR and contain active nucleoli that produce increased rRNA levels. These findings represent a new understanding of at least one way cancer cells resist cytotoxic therapy and contribute to cancer lethality.

We observed dramatic increases in the nuclear volume of cells post-treatment removal over time (**Fig. 2**), representing a 70-fold increase over untreated cells at the latest timepoint measured. Combined with evidence of continued S-phase DNA synthesis in cells that survived after cisplatin treatment, this suggests that surviving cancer cells have undergone multiple whole genome doublings, or polyploidization events. It is generally accepted in the field that polyploid cells that form after chemotherapeutic treatment are multinucleated [4, 10, 11, 26]. We found, however, that not only do mononucleated cells exist, but that a mononucleate morphology is the predominant cell type over the course of cellular recovery from cisplatin, docetaxel, and etoposide therapy in PC3 cells. Nuclear morphology (i.e., mononuclear vs multinuclear) provides valuable insight regarding mechanisms of therapeutic survival. We show that the surviving cells engage in active DNA replication without entry into mitosis (**Fig. 3**), a process known as endocycling, providing a mechanistic explanation for the increase in nuclear size that we observed and that others have observed in studies of similarly large, polyploid cancer cells [4, 6, 10, 11, 26]. Meanwhile, cytoplasmic volumes steadily increased over time, suggesting consistent biosynthesis of other non-genomic cellular contents. With increasing nuclear volume, we further show that treated cells had decreasing nuclear sphericity as a function of recovery time. Nuclear circularity, and its 3D counterpart sphericity, is a representative feature of morphological abnormalities [27, 28]. Nuclear deformability is critical for successful invasion and migration during the metastatic process [5, 17], and decreased sphericity implies capacity for deformability. The decreased sphericity of surviving cells is consistent with previous reports that large, polyploid cancer cells may have increased metastatic potential [29, 30].

To investigate the cellular response to treatment-induced DNA damage in treated cells that eventually die versus those that survive following release from treatment, we simultaneously evaluated DNA damage via gH2AX staining and initiation of DNA damage repair via recruitment of 53BP1. As expected, we initially observed a high level of catastrophic and widespread DNA damage (including increased gH2AX foci counts and gH2AX pan-staining) that decreased as cell populations recovered (**Fig. 5A, B**), consistent with the observed high levels of apoptosis at early timepoints (**Fig. 1C**). Conversely, colocalization of gH2AX and 53BP1 foci as a surrogate for initiation of DDR increased over recovery time and colocalization was increased in mononucleated versus multinucleated cells at every time point (**Fig. 5D, F**). Simultaneously, the relative death rate of treated cells decreased with time (**Fig. 1B**), suggesting that intact DDR in mononucleated cells is associated with survival. Notably, 53BP1 nuclear pan-staining, recognized as a failure of its recruitment to DNA damage sites [22, 31, 32], was not observed at any timepoint, suggesting that DDR was intact in the surviving cell populations.

Together, these data support a paradigm where soon after therapy release, the treated population contains a higher proportion of cells with a high level of widespread and catastrophic DNA damage that leads to apoptosis, while those cells that have successful DDR are more likely to access a pro-survival state. Within the polyploid cell populations that initially survive therapeutic insult, we find that multinucleated cells often exhibit high rates of mitotic catastrophe, even at later timepoints following therapy release (**Fig. 5G, H**). Over recovery time, we observed fewer multinucleated cells in the surviving population, simultaneous with a lower rate of cell death (**Fig. 4**), further indicating that mononucleated cells are more successful at survival once the chemotherapeutic treatment is released. The presence of additional genomic material inside an enlarged, mononucleated cancer cell may reinforce the genetic stability of these cells and enhance their resiliency against varied environmental stressors.

The nucleolus plays a crucial role in nuclear homeostasis as an RNA production hub [23, 24]. We demonstrate that cells that survive long after cisplatin treatment contained fewer nucleoli of a larger size with a corresponding increased rRNA production (**Fig. 6B, C; Fig. S5**). This distinct pattern may be a consequence of endocycling since in normally dividing cells the nucleolar structure disassembles at the mitotic phase and reassembles at interphase [23]. Smaller but more numerous nucleoli are highly correlated with cancer malignancy and have previously been used as a diagnostic signature [25], while our data suggest that fewer, larger nucleoli can also be indicative of cancer cells that persist despite chemotherapeutic insult. Interestingly, stem cells, however, have been reported to have a similar nucleolar signature as surviving cells: fewer but larger nucleoli than differentiated somatic cells [33]. Other studies of large, polyploid cancer cells have suggested that they may act as a type of cancer stem cell, possessing stem cell-like features and cellular programs [4, 6, 8, 9, 11, 15], highlighting the potential of exploring these surviving cells as critical recurrence-mediating cells.

These observations support an underappreciated conserved mechanism of cancer cell survival following stresses such as anti-cancer therapy that has been reported in the literature as the polyaneuploid cancer cell (PACC) state, polyploid giant cancer cells (PGCCs), pleomorphic giant cancer cells, osteoclast-like cells, and persister cells, among other names. The PACC state is initiated when a cancer cell is subjected to a stressor, such as chemotherapy or radiotherapy, triggering a conserved evolutionary program that results in sustained polyploidization of the aneuploid genome (polyaneuploidy) and increased cell size and cell contents. When the cellular stressor is no longer present, and after a period of recovery, cells exit the PACC state and reinitiate mitotic proliferation, a phenomenon clinically observed as cancer recurrence [4, 9–12]. Cells in the PACC state have been identified in the tumors of patients with various cancer types including prostate, ovarian, and breast [34–36], and presence of cells in the PACC state in clinical radical prostatectomy specimens from prostate cancer patients is predictive of an increased risk of disease recurrence [37]. The fundamental characteristics of the PACC state, however, including nuclear morphology (critical for understanding DNA packaging of a multi-polyploid cell) and mechanism of entry and maintenance in this PACC state, are unclear.

Growing investigation into the role of the PACC states relevant to cancer lethality suggests that the PACC state is a “hallmark of lethal cancer” [9]. The findings presented in this study suggest that mononucleated cells in the PACC state are critical players in therapy survival and to eventual cancer recurrence. These data indicate at least two distinct fates of cancer cells following treatment: 1) cells with catastrophic, irreparable DNA damage undergo apoptosis or other forms of cell death; and 2) cells engage DDR survive and engage in continued endocycling, leading to a predominantly mononuclear cell population with increased nuclear size and nucleolar size and capacity. We showed that mononuclear cells in the PACC state had higher evidence of DDR and increased survival, indicating mononucleation, or the processes that lead to that nuclear morphology, offers a survival advantage over multinucleation. Conversely, cells with a multinucleate morphology have apparently lower levels of DDR, higher rates of mitotic catastrophe, and lower survival over time. Given the increased nuclear size and endocycling observed in mononucleated cells in the PACC state, it is likely that these multinucleated cells eventually activate the cell-cycle checkpoints that they initially bypass to enter a polyploid state, thus triggering apoptosis.

While this is a key observation, further work is needed to elucidate if this multinucleation is the cause or consequence of such cell cycle checkpoint activation. The characterization of structural morphological characteristics of the resistant cells in the PACC state highlight a number of compelling routes for future study, including the structural phenotypes that are associated with and / or enable survival of malignant cells. Additional studies are needed to understand the unique and shared characteristics and plasticity of the cells in the PACC state with other cell states that share these structural characteristics, e.g., senescent cells and stem cells. Further characterization of the cellular and molecular programs that cancer cells in this state can be leveraged to understand phenotypes that contribute to cancer lethality.

## Abbreviations

PACC: polyaneuploid cancer cell

## Acknowledgments

The authors thank members of the Cancer Ecology Center for thoughtful discussion and feedback.

## Author contributions

<u>Kim, C-J:</u> conceptualization, methodology, investigation, writing (original draft), writing (review and editing), visualization. <u>Gonye, ALK:</u> conceptualization, methodology, investigation, writing (original draft), writing (review and editing), visualization. <u>Truskowski, K:</u> methodology, investigation. <u>Lee, C-F:</u> investigation.<u>Cho, Y-K:</u> conceptualization, writing (review and editing), funding acquisition. <u>Austin, RH:</u> conceptualization, writing (review). <u>Pienta, KJ:</u> conceptualization, writing (review and editing), funding acquisition. <u>Amend, SR:</u> conceptualization, writing (review and editing), funding acquisition.

## Data availability statement

Data were generated by the authors and available on request from the corresponding authors.

## Supplementary materials

### Materials and methods

#### Nuclei isolation

Cells were lifted, washed twice with PBS, and stored on ice. Nuclei extraction buffer (NEB) was prepared on ice and consisted of final concentrations of 10 mM HEPES (Life Technologies Corp., #15630-080), 1.5 mM MgCl_2_, 10 mM KCl, 0.25% NP-40 (Thermo Scientific, #95124), and 1X protease/phosphatase inhibitor (Thermo Scientific, #1861281) diluted in ultrapure deionized water. 500 μL of NEB was added per 1×10^6^ cells, samples were mixed six times via pipetting and then placed on ice for 10 minutes. Samples were pelleted for 10 seconds in a compact benchtop centrifuge, and the supernatants containing cytoplasmic components were removed. This was repeated twice more, with 5-minute incubations on ice. Following the final removal of supernatants, samples were resuspended in 10% formalin, fixed for 10 minutes, and then filtered through a 70 μm cell strainer. The fixed/filtered nuclei were then pelleted with a quick spin, formalin was removed, and nuclei were resuspended in PBS. Nuclei suspensions were allowed to adhere to 35 mm Ibidi *μ*-Dishes (glass bottom, Ibidi USA Inc., #81158) pre-coated with Poly-L-Lysine (EMD Millipore Corp., #A-005-C) for 30 minutes at room temperature. Once any non-adhered nuclei were washed away, samples were permeabilized with 0.5% Triton-X 100 in PBS and stained for immunofluorescence imaging, then covered with ProLong™ Diamond Antifade Mountant with DAPI. Images were analyzed for nuclear area via a custom CellProfiler 4.0 pipeline.

#### Western blotting

Cells were lysed in ice-cold lysis buffer [150 mmol/L NaCl, 1% Triton X-100, 0.5% sodium deoxycholate, 0.1% SDS, 50 mmol/L Tri (pH 8.0), protease inhibitor cocktail (Roche)], and cell debris was removed by centrifugation at 4 °C for 10 minutes at 12,000 rpm. Equal masses of protein were resolved via SDS-PAGE (gels: Bio-Rad Laboratories, #4561095) and transferred onto Trans-Blot Turbo nitrocellulose membranes (Bio-Rad Laboratories, #1704158). Membranes were blocked with 1X casein blocking buffer (Sigma–Aldrich, #B6429,) for 1 hour at room temperature, then incubated with primary antibodies against P-NPM, NPM, β-Actin, and appropriate secondary antibodies (see **Table S1**). Blots were imaged on a LICOR Odyssey and quantified by ImageJ software. The relative protein expression level in each sample was normalized to the internal control.

#### FUCCI cell line generation

1.5×10^6^ HEK-293T cells were plated in 10 cm dishes in DMEM supplemented with 10% FBS. After 24 hours, cells were transfected for second generation lentivirus packaging as follows: 1 μg of Ki67p-T2A-FUCCI plasmid(67) (gifted from Alexander Zambon) and 1 μg of psPAX2:pMD2.g (packaging:envelope) at an 8:1 ratio were diluted in 200 μL of OptiMEM media (Gibco, #31985062). Both psPAX2 (Addgene plasmid #12260; http://n2t.net/addgene:12260; RRID:Addgene_12260) and pMD2.g (Addgene plasmid #12259; http://n2t.net/addgene:12259; RRID:Addgene_12259) were a gift from Didier Trono. 5 μL of X-tremeGENE HP transfection reagent (Roche, #XTGHP-RO) was added, and the solution was mixed by pipetting and incubated for 30 minutes before being added dropwise onto the HEK-293Ts. After 18-20 hours, media was changed to the target cell media plus 30% heat inactivated FBS. Packaging cells were allowed to condition media for 48 hours before viral harvest. Viral media was removed and filtered through a Millex HP 0.45 μm PES syringe filter (Millipore, #SLHPM33RS). Lentivirus from 1-2 10 cm dishes (10-20 mL) was concentrated using Clontech Lenti-X concentrator (Takara, #631232) per the manufacturer’s protocol. The viral pellet was resuspended in target cell media supplemented with 30% heat inactivated FBS and added onto PC3 cells (3.5×10^5^/well) in 6-well plate along with 10 μg/mL Polybrene Infection Reagent (Millipore, #TR-1003). PC3 cells were transduced for 20 hours before media was changed to fresh supplemented RPMI. After 3 days of recovery, cells were selected with 5 μg/mL blasticidin (Gibco, #A1113903) and expression of reporters was assessed on an EVOS FL Auto imaging system (Life Technologies) with GFP (for mAG) and TxRed (for mCherry) light cubes. Clones were generated from the selected cell pool via limiting dilution cloning.

#### EdU incorporation assay

Large cells were generated as described and then cultured for 1, 3, 5, 9, and 12 days after being filtered following 72 hours of drug treatment. The cells then were seeded on 12 mm round coverslips in 24-well plates. Once cells adhered, the media was replaced and supplemented with 10 μM EdU (Molecular Probes, #E10187). After a 20-hour incubation period under standard conditions, the Click-It Plus EdU assay (Molecular Probes, #C10640) was performed according to the manufacturer’s protocol. Coverslips were mounted to slides using ProLong™ Diamond Antifade Mountant with DAPI (see above). Images were acquired using the 10x objective on the Zeiss Observer Z1 microscope and ZEN pro 2.0 software (Carl Zeiss Microscopy). A custom CellProfiler pipeline was developed and used to quantify total number of nuclei in the tiled image based on DAPI signal and number of EdU-positive nuclei based on AlexaFluor™ 647 signal within nuclei boundaries.

#### Ribosomal RNA gel electrophoresis

Total RNA was isolated from cells using the PureLink™ RNA Mini Kit (Invitrogen, # 12183025). A mass of 100 ng total RNA was prepared for gel electrophoresis and run on a 1% agarose-TBE gel with ethidium bromide. Gels were imaged on a Bio-Rad ChemiDoc™ XRS+ and 28s, 18s, and 5.8s ribosomal RNA bands could be observed.

### Supplementary figures

**Figure S1.**
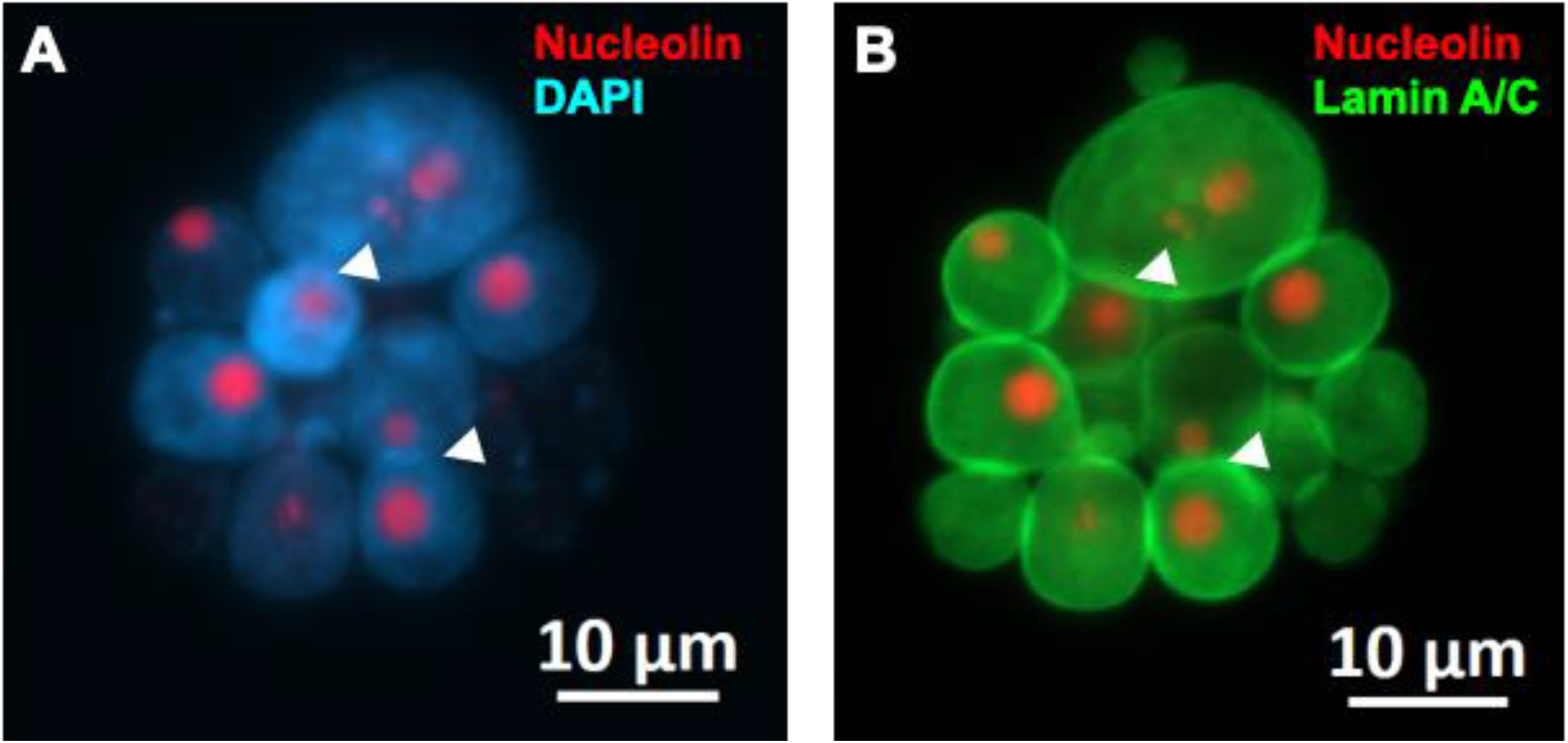
Differences in nuclear visualization between DAPI and lamin A/C. A cisplatin-treated PC3 cancer cell 1 day post-treatment removal was stained for nucleolin and lamin A/C and mounted with DAPI prior to immunofluorescent imaging. Differences in the ability of **(A)**DAPI vs. **(B)**lamin A/C to discriminate individual nuclei should be noted.

**Figure S2.**
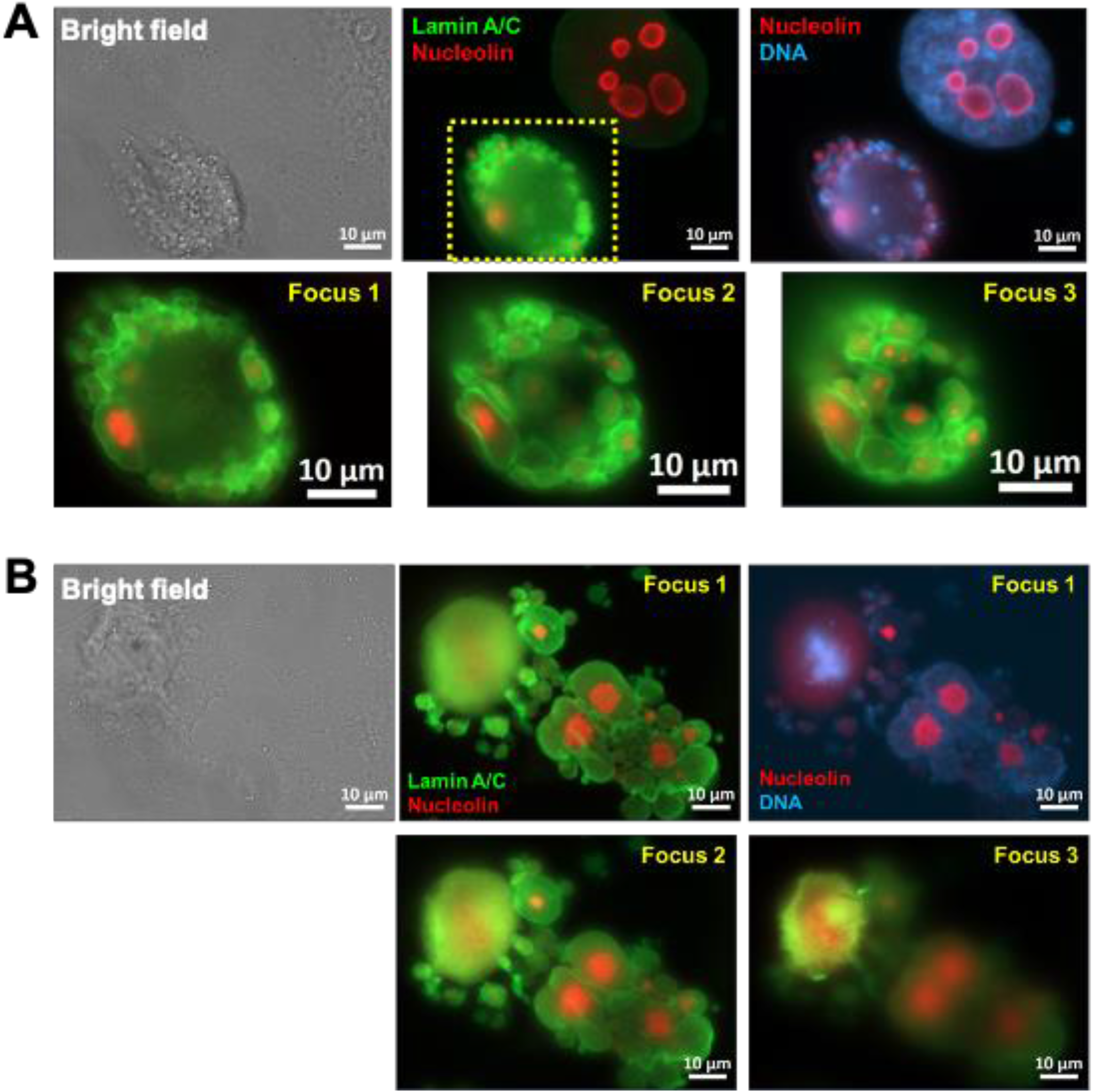
Effects of focal plane on morphologic discrimination of multinucleated cells. Treated PC3s induced with 72h 6 μM cisplatin treatment were stained for lamin A/C, nucleolin, and mounted with DAPI. **(A)**and **(B)**show two different examples of cells where the number of micronuclei/extent of multinucleation might be missed in 2D immunofluorescent microscopy due the effect of different focal planes and cell height.

**Figure S3.**
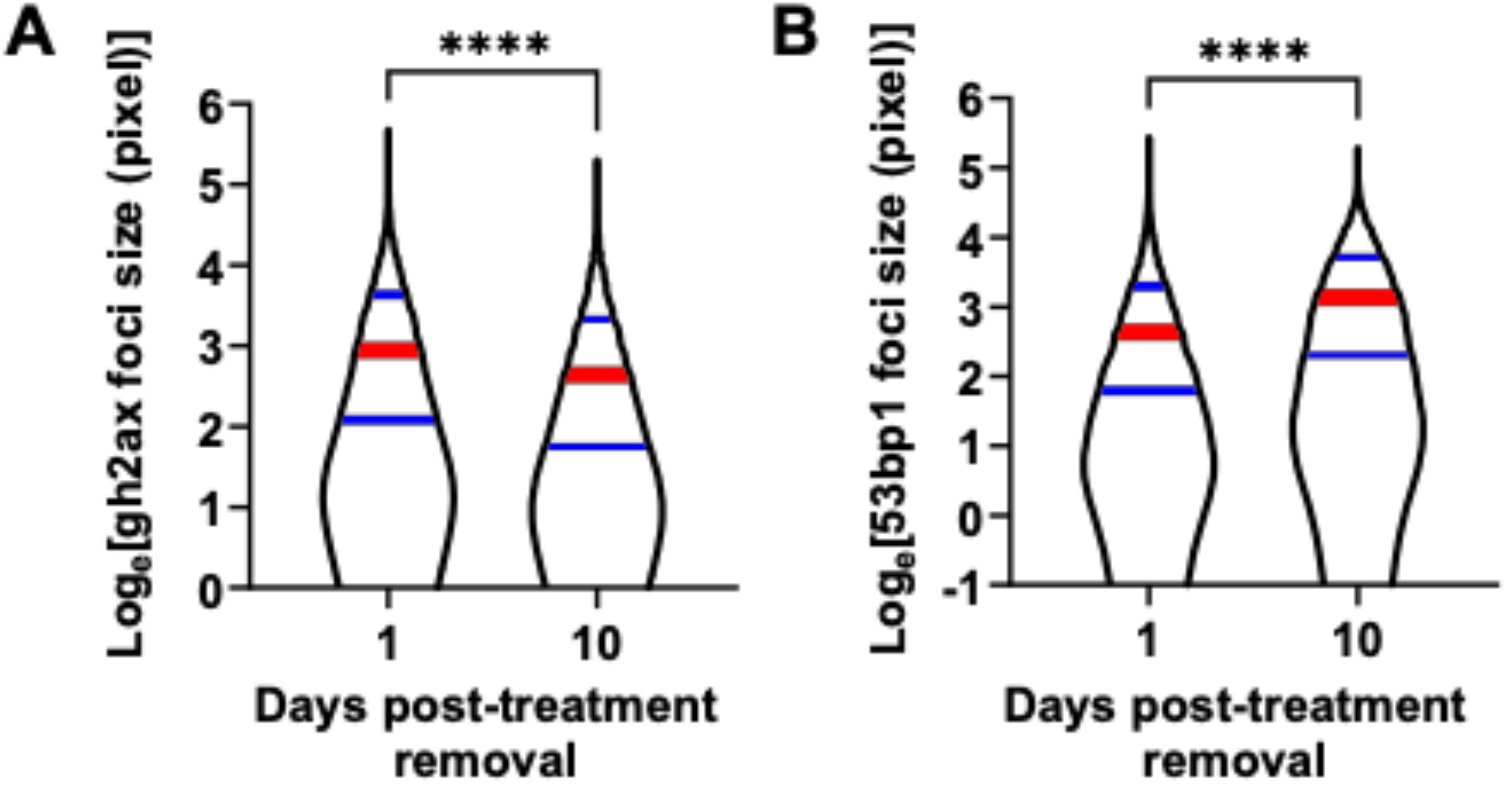
gH2AX and 53BP1 foci size in cisplatin treated PC3s 1 and 10 post-treatment removal. Treated PC3 cells were generated with 72h 6 μM cisplatin treatment and immunofluorescence staining/imaging was used to visualize genomic contents (DAPI), sites of DNA damage (gH2AX), and DDR machinery recruitment (53BP1). A custom CellProfiler 4.0 pipeline was used to determine **(A)**the size of gH2AX foci and **(B)**the size of 53BP1 foci. Mann-Whitney tests were used to detect differences between groups, **** = p<0.0001.

**Figure S4.**
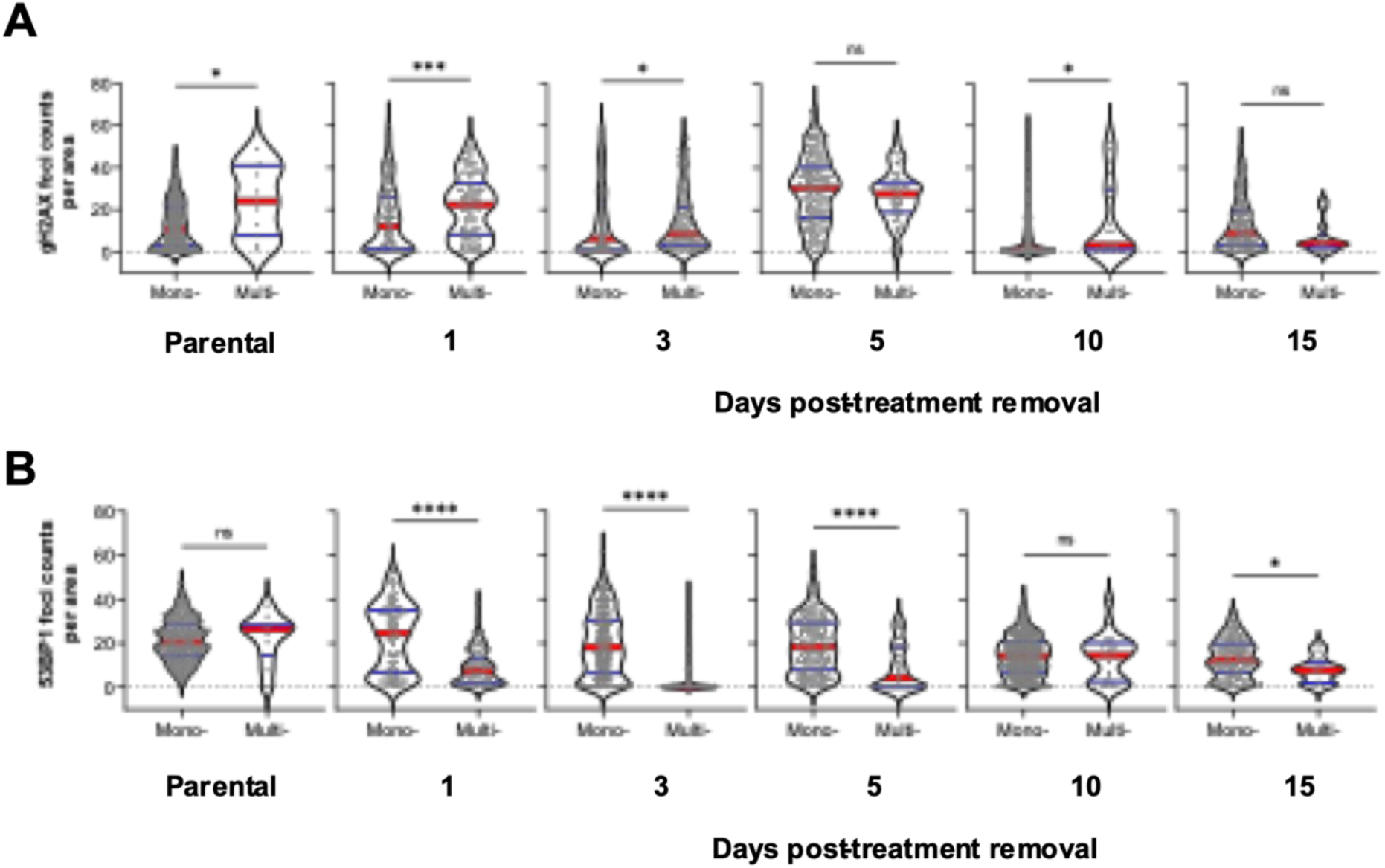
gH2AX and 53BP1 foci counts in cisplatin treated PC3s stratified by nuclear morphology. PC3 cells were treated with 6 μM cisplatin for 72h. Immunofluorescence staining and imaging was used to visualize genomic contents (DAPI), sites of DNA damage (gH2AX), and DDR machinery recruitment (53BP1) within nuclei. For each day post-treatment recovery, cells were classified as mononucleates and multinucleates, and a custom CellProfiler 4.0 pipeline was used to determine raw count per unit cellular area of the number of **(A)**gH2AX foci and **(B)**53BP1 foci.

**Figure S5.**
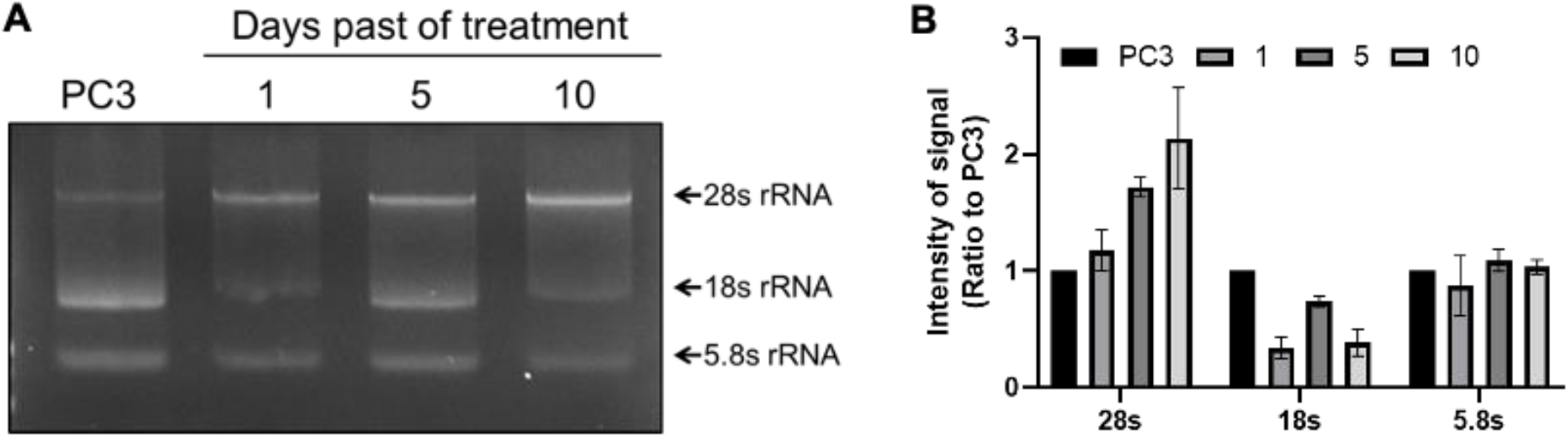
Ribosomal RNA expression levels in treated PC3s. RNA was isolated from treated PC3s induced with 72h 6 μM cisplatin at various points following treatment removal. **(A)**100 ng total RNA was resolved via gel electrophoresis, and the rRNA pattern (28s, 18s and 5.8s rRNA) signals was observed via staining with EtBr. **(B)**Densitometry was used to quantitate differences in rRNA levels from three independent replicates.

**Supplementary Table S1.** Antibodies used in this study.

| PRIMARY ANTIBODIES |  |  |  |  |
| --- | --- | --- | --- | --- |
| Antigen | Species | Dilution | Source |  |
| lamin A/C and | Mouse IgG2a | 1:100 (IF) | Cell Signaling Technologies, #4777 |  |
| Alexa Fluor<sup>TM</sup> 647-conjugated nucleolin | Rabbit | 1:50 (IF) | Abcam, #ab202709 |  |
| gamma H2A.X | Mouse IgG1 | 1:250 (IF) | Abcam, #ab22551 |  |
| 53BP1 | Rabbit | 1:100 (IF) | Abcam, #ab175933 |  |
| NPM | Rabbit | 1:1000 (WB) | Cell Signaling Technologies, #3542 |  |
| Phospho-NPM | Rabbit | 1:1000 (WB) | Cell Signaling Technologies, #3541 |  |
| β-Actin | Mouse | 1:10000 (WB) | Millipore Sigma, #A5441 |  |
| SECONDARY ANTIBODIES |  |  |  |  |
| Conjugate | Host species | IgG species | Dilution | Source |
| Alexa Fluor 488 | Goat | Mouse IgG1 | 1:1000 (IF) | Invitrogen, #A21121 |
| Alexa Fluor 594 | Goat | Rabbit | 1:1000 (IF) | Invitrogen, #A32740 |
| Alexa Fluor 647 | Goat | Mouse IgG2a | 1:500 (IF) | Invitrogen, #A21241 |
| IRDye 680RD | Goat | Mouse | 1:20000<br>(WB) | LI-COR Biosciences,<br>#926-68070 |
| IRDye 800CW | Goat | Rabbit | 1:15000<br>(WB) | LI-COR Biosciences,<br>#926-32211 |

